# Discovery and characterization of *Alu* repeat sequences via precise local read assembly

**DOI:** 10.1101/014977

**Authors:** Julia H Wildschutte, Alayna Baron, Nicolette M Diroff, Jeffrey M Kidd

## Abstract

*Alu* insertions have contributed to >11% of the human genome and ~30–35 *Alu* subfamilies remain actively mobile, yet the characterization of polymorphic *Alu* insertions from short-read data remains a challenge. We build on existing computational methods to combine *Alu* detection and *de novo* assembly of WGS data as a means to reconstruct the full sequence of insertion events from Illumina paired end reads. Comparison with published calls obtained using PacBio long-reads indicates a false discovery rate below 5%, at the cost of reduced sensitivity due to the colocation of reference and non-reference repeats. We generate a highly accurate call set of 1,614 completely assembled *Alu* variants from 53 samples from the Human Genome Diversity Project panel. We utilize the reconstructed alternative insertion haplotypes to genotype 1,010 fully assembled insertions, obtaining >99% agreement with genotypes obtained by PCR. In our assembled sequences, we find evidence of premature insertion mechanisms and observe 5’ truncation in 16% of *Alu*Ya5 and *Alu*Yb8 insertions. The sites of truncation coincide with stem-loop structures and SRP9/14 binding sites in the *Alu* RNA, implicating L1 ORF2p pausing in the generation of 5’ truncations. Additionally, we identified variable *Alu*J and *Alu*S elements that likely arose due to non-retrotransposition mechanisms.

## INTRODUCTION

Mobile elements (MEs) are discrete fragments of nuclear DNA that are capable of copied movement to other chromosomal locations within the genome (1). In humans, the ~300 bp *Alu* retroelements are the most successful and ubiquitous MEs, collectively amounting to >1.1 million genome copies and accounting for >11% of the nuclear genome (2,3). The vast majority of *Alu* insertions represent events that occurred in the germline or early during embryogenesis (4) millions of years ago and now exist as non-functional elements that are highly mutated and no longer capable of mobilization (3). However, subsets of MEs, including *Alu* and its autonomous partner *L1Hs,* remain active and continue to contribute to new ME insertions (MEIs), resulting in genomic variation between individuals (5) and between somatic tissues within an individual (6,7).

The human genome contains elements derived from the *Alu*Y*, Alu*S, and *Alu*J lineages, which can be further stratified into more than ~35 subfamilies based on sequence diversity and diagnostic mutations (2,5,8). Most human *Alu* elements are from the youngest lineage, *Alu*Y, whose members have been most actively mobilized during primate evolution (5,9). Of these, the *Alu*Ya5 and *Alu*Yb8 subfamilies have contributed to the bulk of insertions in humans (10–14), although polymorphic insertions from >20 other *Alu*Y and >6 *Alu*S subfamilies have also been reported (5,15), implying polymorphic insertions of other lineages may still be segregating. In contemporary humans, the retrotransposition of active *Alu* copies results in *de novo* germline insertions at a frequency of ~1:20 live births (11,16). Over 60 novel *Alu* insertions have been shown to cause mutations leading to disease (17–19); older, existing *Alu* insertions in the genome have also been shown to facilitate the formation of subsequent rearrangements by providing a template of sequence utilized in non-allelic homologous recombination, replication-template switching, and the repair of double-strand breaks (2,18–26). Thus, *Alu* insertions continue to shape the genomic landscape and are recognized as profound mediators of genomic structural variation.

Active copies of *Alu* are non-autonomous but contain an internal RNA Pol III promoter (27). Mediated by L1 encoded enzymes, *Alu* transcripts are mobilized by a ‘copy-and-paste’ mechanism referred to as target primed reverse transcription (TPRT) (7,28). TPRT involves the reverse transcription of a single stranded *Alu* RNA to a double stranded DNA copy, during which two staggered single-stranded breaks are introduced in the target DNA of ~5 to ~25 bp that are later filled by cellular machinery. The resulting structure consists of a new *Alu* flanked by characteristic target site duplications (TSDs) and a poly-A tail of variable length. Together these serve as hallmarks of retrotransposition. Integration of the new copy is permanent, although *Alu* can be removed by otherwise encompassing deletions, or very rare recombination-mediated excision events (29). Complete TPRT of a full-length element is responsible for the majority of *Alu* insertions. A minority of insertions have signatures of 5’ truncation but otherwise appear as standard retrotransposition events, indicating their movement by premature TPRT (21,30,31).

The primary difficulty in identifying novel *Alu* insertion loci stems from the abundant copy-number of these elements in primate genomes. Various approaches for large-scale analyses of *Alu* and other ME types have been developed that utilize next generation sequencing platforms. Scaled sequencing of targeted *Alu* junction libraries has permitted genome-wide detection, as implemented in techniques such as Transposon-Seq (32) and ME-Scan (33,34). Such targeted methods offer high specificity and sensitivity, but are restricted by the primers used for detection and are generally subfamily-specific. A broader detection of *Alu* variant locations is possible by using computational methods to search Illumina whole genome sequence (WGS) paired reads by ‘anchored’ mapping. This method seeks to identify discordant read pairs where one read maps uniquely to the reference (*i.e.,* the ‘anchor’) and its mate maps to the element type in query (10,35,36). However, in considering read pair data alone, in their simplest form these methods are limited in the recovery of variant-genome junctions that might otherwise be captured from split read information. Specialized algorithms now offer improved breakpoint accuracy from WGS data by consideration of split-reads, soft-clipped reads, and unmapped reads during variant detection (36–39). Beyond these basic requirements, existing programs differ in the implementation of additional filters and read-support criteria to identify the subset of calls likely to be true. For most callers this includes removing MEI candidate calls that are located near reference MEs of the same class due to their higher likelihood of being false predictions ((36,38–41) but see (37)). Thus, as in all methods, a trade-off exists between sensitivity and specificity.

Having been mostly applied to high-coverage WGS, these methods require modification when applied to lower coverage data. Generally, these read-based detection methods have been developed in the context of ME variant loci discovery and reporting. Assembly approaches have received increasing application to next-generation sequencing data for breakpoint identification. Tools such as TIGRA (42), a modification of the SGA assembler used in HYDRA-MULTI (43,44), and the use of a *de brujin* graph-based approach in SVMerge (45) have been developed to assemble structural variant breakpoints from population scale or heterogeneous tumor sequencing studies. Assembly-based approaches have also lead to increased sensitivity and specificity for the detection of SNPs and small indels (46–50) and mobile elements (38). The sequence of polymorphic *Alus* has also been inferred by remapping supporting reads to an element reference followed by the construction of a consensus sequence (14). Since sequence changes within elements can determine their activity (9), the assembly of insertion sequences can better inform our understanding of element proliferation. Additionally, insertion haplotype reconstruction offers a direct way to determine genotypes across samples based on an analysis of sequence data supporting each allele.

Here, we utilize a classic overlap-layout-consensus assembly strategy applied to ME-insertion supporting reads to completely reconstruct and characterize *Alu* insertions. We apply this approach to pooled WGS data, from 53 individuals in 7 geographically diverse populations from the Human Genome Diversity Project (HGDP) panel (51,52). We assess the limitations of this approach by cross-comparison with *Alu* insertions identified from PacBio long sequencing reads in a complete hydatidiform mole (CHM1) (53). Finally we demonstrate the ability to obtain accurate genotypes based on explicit mapping to reconstructed reference and alternative alleles. We present the analysis of 1,614 fully reconstituted *Alu* insertions from these samples, including breakpoint refinement and genotyping of 1,010 insertions. These results provide a basis for future study of such MEIs in human disease and population variation, and should facilitate similar analyses in relevant non-human models.

## MATERIAL AND METHODS

### Samples

We analysed whole genome, 2x101 bp Illumina read sequence data from a subset of 53 samples and 7 populations from the HGDP: Cambodia (HGDP00711, HGDP00712, HGDP00713, HGDP00715, HGDP00716, HGDP00719, HGDP00720, HGDP00721), Pathan (HGDP00213, HGDP00222, HGDP00232, HGDP00237, HGDP00239, HGDP00243, HGDP00247, HGDP00258), Yakut (HGDP00948, HGDP00950, HGDP00955, HGDP00959, HGDP00960, HGDP00963, HGDP00964, HGDP00967), Maya (HGDP00854, HGDP00855, HGDP00856, HGDP00857, HGDP00858, HGDP00860, HGDP00868, HGDP00877), Mbuti Pygmy (HGDP00449, HGDP00456, HGDP00462, HGDP00471, HGDP00474, HGDP00476, HGDP01081), Mozabite (HGDP01258, HGDP01259, HGDP01262, HGDP01264, HGDP01267, HGDP01274, HGDP01275, HGDP01277), and San (HGDP00987, HGDP00991, HGDP00992, HGDP01029, HGDP01032, HGDP01036). WGS data was processed using BWA, GATK (54) and Picard (http://picard.sourceforge.net) as described previously (52) and is available at the Sequence Read Archive under accession SRP036155. Final datasets are ~7x coverage per sample. For analysis of CHM1 we utilized Illumina data obtained under accession SRX652547, and ERX009608 for analysis of NA18506. Reads were mapped respectively to the hg19/GrCh37 or hg18/build36 genomes as appropriate using same procedures described above.

### Non-reference *Alu* discovery

RetroSeq ‘discover’ was run on individual BAM files for each sample to identify discordant read pairs with one read mapping uniquely to the reference genome and its pair to an *Alu* consensus or to an annotated *Alu* present in the reference. A FASTA file of *Alu* consensus sequences was obtained from RepBase (version 18.04) (55). Reference *Alu* elements were excluded using hg19 RepeatMasker (56) annotations. Candidate loci were assessed using RetroSeq ‘call’. For this analysis, we combined the supporting read information discovered in each individual and ran the ‘call’ phase on a combined BAM consisting of all samples. A minimum of 2 supporting read pairs was required per call (-reads). A maximum read-depth of 1000 (-depth; default is 200) was utilized for regions surrounding each call in order to accommodate the increased coverage of the merged BAM. Finally, any output call within 500 bp of an annotated *Alu* insertion was excluded using the bedtools window command (57) and RepeatMasker hg19 reference annotations (56). Unless otherwise noted, any other RetroSeq options were run at the default settings. Final *Alu* calls having met the further criteria of a filter tag FL=6,7,or 8 were selected for subsequent analysis.

### Assembly of non-reference *Alu* elements

*De novo* assembly of insertion-supporting reads for each candidate insertion was performed using CAP3 (58). For each candidate insertion, a window of 200 bp was defined around the predicted breakpoint, from which we extracted read-pairs reported to support an insertion at that site based on RetroSeq outputs, and intersecting read-pairs with a soft-clipped segment from the BAM corresponding to each sample. For soft-clipped reads we required the clipped portion to be ≥ 20 bp with a mean quality ≥ 20. Using CAP3 we then assembled the extracted reads per site, with parameters chosen to account for shorter matches (-c 25 -j 31 -o 16 -s 251 -z 1 -c 10). CAP3 utilizes read-pair information to report scaffolds of contigs that are linked together but without assembled overlap. We merged such contigs together, separated by 300 ‘N’ characters to represent sequence gaps in the assembly. The resulting contigs and scaffolds were analysed using RepeatMasker (56), from which 2,971 candidate assembled sites were identified that contain an *Alu* element (≥ 30 bp match) and at least 30 bp of flanking non-gap sequence. This pipeline is available on GitHub at https://github.com/KiddLab/insertion-assembly.

### Breakpoint determination

Breakpoints for the assembled *Alu* variants were recovered utilizing a multiple alignment-based approach similar to an approach previously described (59,60). Orientation of candidate insertion sequences relative to the reference genome was determined using BLAT (61). Candidate breakpoints were then identified using *miropeats* (62) followed by a semi-automated parsing process. A global alignment was obtained for sequences from the two insertion breakpoints to the corresponding segment on the reference genome using *stretcher* (63) with default parameters, to generate pairwise alignments for two sequences aligned independently against the third (*i.e.,* reference) sequence. A 3-way alignment was created from the two pair-wise alignments by inserting gaps as appropriate. Alignment columns were scored as a match among all three sequences (‘*’), a match between the left (‘1’) or right (‘2’) insertion breakpoint and the reference, or mismatch among all sequences (‘N’). We then computed an alignment score across the left and right breakpoint sequence, with matches between the target sequence and the genome sequence (‘1’ or ‘*’ for the left breakpoint and ‘2’ or ‘*’ for the right breakpoint) resulting in a score of +1, a sequence mismatch among all three sequences a score of −1, and a match among the reference and the other breakpoint a score of −3. The same procedure was applied to the right breakpoint, except the score was tabulated from right to left across the 3-way alignment. Visualizations of the resulting aligned sequences with breakpoint annotations were constructed and subjected to manual review. When necessary, the sequences extracted for breakpoint alignment were adjusted and the alignment and scoring scheme described above then repeated until a final curated set of 1,614 assembled insertions was obtained. The breakpoint was then interpreted as the position where the maximum cumulative score was reached respective to the reference, from which regions of overlapping sequence on the reference allele were determined. Of note, 1) overlapping regions are defined from the alignment itself, without regard to the element boundaries, and 2) in scoring, a small degree of divergence among regions of overlap is permitted, in some cases, resulting in the identification of longer segments with less than 100% identity. To define candidate TSDs, segments of identical sequence were identified within the region of sequence overlap by extending toward the element 3’ end until a mismatch was observed.

### Sub-family assignment and analysis

A multiple sequence alignment was constructed from 45 *Alu* consensus sequences corresponding to active elements reported by (5) and all subfamilies identified in the assembled set by RepeatMasker. Consensus sequences were obtained from RepBase (version 18.04) (55) and aligned using MUSCLE v3.8.31 (64) run with default parameters. Each of the of 1,614 assembled *Alu* sequences was extracted and separately aligned to the consensus profile alignment. The proportion of sequence differences between each element and each sub-family consensus was tabulated, excluding the poly-A tail. Elements equally distant from multiple subfamilies were deemed to be unclassified. Alignments for the 1,010 sites suitable for genotyping were utilized to assess the extent of the recovered element length relative to the subfamily consensus.

### Validation

A total of 66 assembled *Alu* insertions were validated by Sanger sequencing. This includes 20 sites selected randomly using a custom python script, 33 sites chosen based on characteristics of the assembled insertions, and 13 sites representing members of *Alu*S and *Alu*J lineages. Chromosomal coordinates for each insertion were considered based on unique mapping of CAP3 assembled contigs and subsequent breakpoint analysis to the hg19 build. We extracted ~500 bp in either direction of each insertion from the hg19 reference (http://genome.ucsc.edu/). The sequence was masked using RepeatMasker, and primers designed to include at least 150 bp in either direction of the predicted insertion, avoiding masked sequence when possible. Each primer set was analysed by *in silico* PCR to the hg19 reference to ensure site-specific target amplification predictions overlapping each breakpoint, and to infer product size predictions for either allele. All primers were designed using Primer3v.0.4.0 (65) and purchased from IDT. Loci examined, primers, and samples analysed are summarized in Table S2.

All PCRs were performed with ~50ng of genomic DNA as template along with 1.5–2.5 μM Mg^++^, 200μM dNTPs, 0.2 μM each primer, and 2.5 U Platinum Taq Polymerase (Invitrogen). Reactions were run under conditions of 2 min denaturation at 95 °C; 35 cycles of [95 °C 30 sec, 55 °C to 59 °C 30 sec, 72 °C 2 min]; and a final extension at 72°C for 10 min. For each PCR reaction, 10uL were analysed by electrophoresis in 1% agarose in 1 × TBE. Products from at least one positive reaction per locus were sequenced. When possible, PCR products from a homozygous individual were sequenced; otherwise the insertion-supporting fragment was gel-extracted (Qiagen), and the products eluted in water and subjected to sequencing. Sequencing was performed with primers situated both upstream and downstream of the insertion to account for uncertainty introduced from polymerase slippage at the poly-A tails and ensure amplification across both predicted breakpoints. Traces obtained for each insertion allele were aligned to the corresponding reference allele and CAP3 assembled contig in order to confirm the presence of the *Alu* insertion, breakpoints, and agreement in nucleotide sequence between the validated and assembled insertion.

### Comparison with previous studies

For comparison with Chaisson *et al.* calls on CHM1 (53), we utilized insertion positions based on hg19 coordinates from http://eichlerlab.gs.washington.edu/publications/chm1-structural-variation/. These data consisted of 1,254 total calls classified as “AluYsimple”, “AluSsimple”, “AluSTR”, or “AluMosaic”. Overlapping calls were counted from intersection of any call located within 100 bp. The 1,727 *Alu* insertion calls for NA18506 were based on alu-detect analysis (37) of ERX009608 as obtained from the Sequence Read Archive. Consistent with that study, calls were relative to the hg18 genome assembly, and any call located within 100 bp was counted as an intersecting site.

### Comparison with primate data

The reference and insertion alleles for each *Alu*S or *Alu*J element were searched against primate reference genomes using BLAT (61) on the UCSC Genome Browser (http://genome.ucsc.edu/) (66) Insertion coordinates were converted to hg18 coordinates using liftOver and compared with polymorphic insertions reported in (67). Intersections within 500 bp were reported.

### Construction of random insertion site distributions

We sampled 1,614 insertion positions throughout the genome matching the known sequence preferences of the L1 endonuclease. To reflect the known L1 EN sequence preferences, we constructed a position probability matrix (PPM) using 99 5-bp L1 EN nic-site sequences reported in Gilbert *et al.* (68), with a pseudocount of 1 added to each column to permit the sampling of unobserved sequences. To create each randomly placed insertion we first randomly (uniformly) selected a position in the reference genome sequence and an insertion strand (+ or −). We then extracted the corresponding 5-bp nic-site, calculated its probability *P* using the computed PPM, sampled a random number *p* (from a uniform distribution) and accepted the site if *p* < P. This process was repeated until 1,614 sites were accepted, to generate one set of randomly sampled insertion positions and then repeated to generate 200 sets of randomly sampled insertions. For some analysis we imposed an exclusion mask that mirrored our discovery criteria such that positions within 500 bp of an *Alu* element in the reference genome were not accepted. Scripts for randomly sampling positions based on a PPM are available at https://github.com/KiddLab/random-sample-by-ppm.

### Genotyping

We performed *in silico* genotyping by mapping relevant reads to a representation of the complete insertion and reference alleles for each site. The reference allele consisted of 600 bp of sequence upstream and downstream of the start and end of any inferred TSD extracted from the hg19 reference. Based on the aligned breakpoints, insertion alleles were created by replacing the appropriate portion of this sequence with insertion sequence, accounting for inferred TSDs or target site deletions. For each site, these insertion and reference alleles constituted the target genome for mapping of reads. A BWA index was constructed from each (bwa version 0.5.9). Mapping and analysis was performed separately for each sample and each site. We extracted read-pairs with at least one read having an original mapping within the coordinates of the targeted reference allele with a MAPQ ≥ 20. The extracted read-pairs were then aligned to the site reference and alternative sequences using bwa aln and bwa sampe (version 0.5.9). We then calculated genotype likelihoods based on the number of read pairs mapping to the insertion or reference alleles, considering the resulting MAPQ values as error probabilities as previously described (69). Read-pairs with equal mappings between reference and insertion sequences have a MAPQ of 0 and do not contribute.

Genotypes were obtained from the resulting raw genotype likelihoods using one of two approaches. For sites on the autosomes and the pseudoautosomal region of the X chromosome, genotype likelihoods for *Alu* insertions were processed, along with previously calculated SNP genotypes using LD-aware refinement using Beagle 3.3.2 (with options maxlr=5000, niteration=10, nsamples=30, maxwindow=2000) (70–72). For sites on the X chromosome, genotypes were obtained using a ploidy-aware expectation-maximization (EM) algorithm that utilized the genotype likelihoods and assumes Hardy-Weinberg Equilibrium across all 53 samples. Briefly, we followed (69) to estimate the allele frequency for each site via EM. Using the estimated allele frequency, we then determined genotype prior probabilities for X-linked alleles assuming Hardy-Weinberg equilibrium. These genotype priors were then combined with the already computed genotype likelihoods to identify the sample genotype with the highest posterior probability. Principal component analysis was performed on the resulting autosomal genotypes using the smartpca program from the EIGENSOFT package (73).

### Genotype Validation

In order to validate *in silico* genotyping and permit estimation of genotyping accuracy, a subset of 11 insertion loci were screened from a panel of ten individuals utilizing gel band assays, for a total of 110 predicted genotypes (Table S6). Locus-specific primer sets flanking each insertion locus were designed as above (see Validation) and utilized for PCR amplification of each sample. All reactions were performed with 50 ng genomic DNA, in cycling conditions of 2min at 95°C; 35× [95°C 30 sec, 55°C–59°C 30 sec, 72°C 1 min], and a final 72°C extension of 3 min. 10uL were analysed by electrophoresis in 1.2% agarose in 1×TBE, and results interpreted by banding patterns that supported either the unoccupied or insertion allele, as based on predicted band sizes from *in silico* PCR and size information for the assembled insertion at that site.

## RESULTS

### Precise assembly of full-length *Alu* variants using read data

To generate an accurate and highly specific collection of non-reference polymorphic *Alu* variants from population-scale WGS data we combined methods utilizing read-based discovery of all possible insertion sites with *de novo* local assembly of supporting reads (Figure 1). We utilized WGS data from a subset of the HGDP collection, specifically consisting of 2×101 bp paired-end libraries from 53 individuals across seven populations exhibiting a cline of diversity reflecting the major migration of humans out of Africa, with a median coverage of ~7x per genome. Given the coverage levels, we anticipated insertions that were private to a single individual were likely to be missed. However, we reasoned that borrowing read information across samples would increase our ability to detect rarer insertions that were nonetheless present in multiple samples, and pooled the data into a single merged BAM from all individuals for an effective coverage of ~429x. Candidate *Alu* insertions were then identified by applying RetroSeq (36) to the merged BAM. This particular program implements standard approaches to identity MEI-supporting read signatures, with performance characteristics that are comparable to other existing callers (for example in (36,39,40)) and assigns a quality score to each call based on the number and mapping characteristics of supporting reads (Figure 1, Methods). Based on these outputs we considered calls with a score of 6 or higher for further filtering. To minimize false calls associated with reference elements, we removed any candidate call that mapped within 500 bp of an annotated *Alu* in the human reference. After exclusion, this resulted in 41,365 putative *Alu* insertions with an assigned quality score 6 or higher.

**Figure 1.**
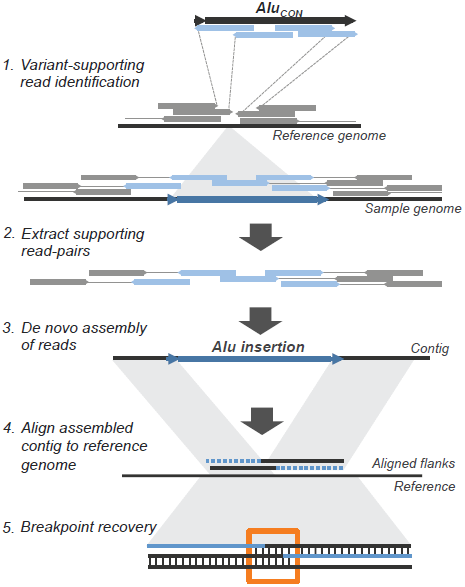
Strategy for detection and assembly of non-reference *Alu* insertions. Approach for reconstruction of non-reference *Alu* insertions from WGS data. 1). WGS in aligned BAM format from 53 samples were merged to a single BAM file, and clusters of *Alu*-supporting read pairs identified using the RetroSeq program by Keane *et al.* 2). *Alu*-supporting read pairs and intersecting split reads were extracted for each candidate site, and 3). Subjected to a *de novo* assembly using the CAP3 overlap-layout assembler 4). *Alu*-containing contigs were then mapped to the reference genome to verify chromosomal coordinates and uniqueness of the call. 5). Breakpoints and putative TSDs from each contig were computationally predicted by 3-way alignment to determine overlap of the assembled upstream and downstream flanks with the pre-insertion site from the hg19 reference.

We then attempted to reconstruct as many individual insertion variants as possible, including the complete *Alu* sequence, its breakpoints, and contiguous flanking sequence for each site. While recent efforts in short read assembly have focused on a *de brujin* graph approach (74–76), we reasoned a local assembly using an overlap-layout-consensus approach would take full advantage of our data. For these purposes we utilized the program CAP3 (58) that was originally developed for the assembly of large-insert clones sequenced using capillary sequencing, but has also been applied to *de novo* assembly of short read RNA-seq (77) and metagenomic sequence data (78). For each putative site, we retrieved any insertion-supporting read pairs as reported by RetroSeq and proximal soft-clipped reads, and then performed *de novo* assemblies with CAP3 (Figure 1) run with parameters adjusted for joining smaller overlaps that could be expected from 101 bp reads (also see Methods). The resulting scaffolds were filtered to require the presence of ≥ 30 bp of *Alu* sequence (with ≥ 90% nucleotide identity) and recovery of ≥ 30 bp of flanking non-gap sequence at one end, resulting in 2,971 candidate assemblies. Assembled scaffolds having satisfied all the above criteria were interpreted to represent the presence of a true non-reference insertion.

The resulting assemblies were aligned to the reference genome and putative breakpoints identified using a semi-automated procedure supplemented by manual curating. For these purposes, we adapted a procedure previously developed for the analysis of structural variant breakpoints represented in finished fosmid clone sequences (59) (also see Methods). A total of 1,614 *Alu-*containing contigs were reconstructed, each having the complete associated insertion and at least one breakpoint with ≥ 30 bp of mapped flanking sequence (Table S1a, Figure S1). Utilizing the locally assembled scaffolds, we determined regions of overlapping sequence on the reference allele (also see Methods; Figure S2). We then identified a set of 1,010 of the 1,614 *Alu* insertions that had both breakpoints flanked by at least 100 bp of non-gap assembled sequences. Given the quality of these assembled calls, we refined the initial regions of overlapping sequence to identify candidate TSDs based on segments of exact nucleotide matches (Figure S3). These 1,010 insertions were deemed to be of the highest quality and suitable for *in silico* genotyping (see below), and had sizes ranging from 77 bp – 495 bp (median 315 bp) with refined candidate TSDs up to 50 bp (median 14 bp).

### Sensitivity and specificity of insertion discovery using short-read assembly

A comprehensive assessment of the performance of MEI callers is hindered by the lack of an orthogonal “gold standard” call set for formal comparison. To better assess the potential limitations of *Alu* discovery using Illumina (2×100 bp) paired-end short reads, we applied our approach to two additional samples, which have been extensively characterized (Table S1b). The first comparison utilized Illumina WGS data generated from a complete hydatidiform mole (CHM1) (53). This sample offers particular advantage for these purposes, as it is essentially haploid throughout its genome and has been subjected to extensive characterization using long PacBio sequencing reads (mean 5.8 kbp). In their analysis, Chaisson *et al.* reported 1,254 *Alu* insertions from this sample; of which 911 intersected within 100 bp of a candidate call based on the Illumina sequence data (Table 1). However, we note these raw RetroSeq calls include 18,501 predictions, implying an extremely high false discovery rate (FDR). Considering only the highest level of RetroSeq support still implies an FDR greater than 80%.

**Table 1.** Sensitivity and Specificity Analysis.

| A. All Calls |  |  |  |  |  |  |  |
| --- | --- | --- | --- | --- | --- | --- | --- |
| Call Set | Predicted Insertions | Overlap with Chaisson et al. | Total Chaisson et al. Sites | Chaisson et al. Sites With Overlap | Sensitivity | Precision | False Discovery Rate |
| RetroSeq Calls | 18,501 | 911 | 1,254 | 896 | 71.5% | 4.9% | 95.1% |
| Support Level >= 6 | 12,721 | 908 | 1,254 | 893 | 71.2% | 7.1% | 92.9% |
| Support Level 8 | 5,294 | 889 | 1,254 | 874 | 69.7% | 16.8% | 83.2% |
| Assembled | 468 | 449 | 1,254 | 449 | 35.8% | 95.9% | 4.1% |

| B. Calls at least 500bp distant from reference Alus |  |  |  |  |  |  |  |
| --- | --- | --- | --- | --- | --- | --- | --- |
| Call Set | Predicted Insertions | Overlap with Chaisson et al. | Total Chaisson et al. Sites | Chaisson et al. Sites With Overlap | Sensitivity | Precision | False Discovery Rate |
| RetroSeq Calls | 924 | 479 | 578 | 473 | 81.8% | 51.8% | 48.2% |
| Support Level >= 6 | 615 | 479 | 578 | 473 | 81.8% | 77.9% | 22.1% |
| Support Level 8 | 520 | 474 | 578 | 468 | 81.0% | 91.2% | 8.8% |
| Assembled | 468 | 446 | 578 | 446 | 77.2% | 95.3% | 4.7% |

Mapping into highly repetitive regions using shorter sequencing reads results in an unacceptably high degree of false calls and necessitates filtering out regions near reference elements, a step common to most mobile element callers (36,38,39,41). The longer reads available from the CHM1 sample permitted interrogation of genome intervals harbouring a high number of repetitive elements. In fact, 54% of *Alu* insertion calls by Chaisson *et al.* are located within 500 bp of an *Alu* sequence present in the human reference genome. Limiting the analysis to only those calls ≥500 bp from any reference *Alu* results in an increase in both sensitivity and precision. Requiring successful element assembly further increases the precision: 446 of 468 of our assembled calls intersect (within 100 bp) with CHM1 reported calls. Assuming the remaining calls are all errors implies a FDR below 5%, however we note additional analysis suggests this may be an over estimate (see Discussion).

To further investigate the precision of our assembly approach, we applied this approach to 2×100 bp Illumina WGS data from the Yoruba sample NA18506, an individual that has been analysed using multiple MEI callers (35,37,38). Again excluding calls within 500 bp of an annotated insertion, initial filtering of RetroSeq calls on the NA18506 sample resulted in 1,375 putative *Alu* insertions having a quality score of 6 or higher. A total of 820 *Alu* insertions were fully assembled, of which 774 intersect with calls reported by alu-detect (out of 1,727 total calls) (37), again implying a FDR around 5%.

### Validation of assembled HGDP insertions

In validation, our specific goals were to demonstrate 1) the presence of the *Alu* at that chromosomal location, 2) agreement between the assembled sequence with the cognate validated sequence for the insertion, and 3) contiguous sequence of each insertion with its mapped flanking regions. To assess the accuracy of our approach, we experimentally validated a set of 20 insertions that were randomly selected from our total set of 1,614, without bias to frequency or queried sample. We further validated 33 sites with unusual breakpoint characteristics and 13 sites derived from the *Alu*S or *Alu*J lineages (Tables S2 and S3). Detailed alignments corresponding to individual insertions, including visualized trace information for all sites, are provided in Figures S4-S6 and properties for all validated sites are summarized in Table S3. Examples of three representative insertions are highlighted in Figure 2, illustrating the recovery of mapped *Alu*-containing contigs, breakpoint estimations at those sites, and alignment of the deduced nucleotide sequence to the corresponding CAP3 assembly.

**Figure 2.**
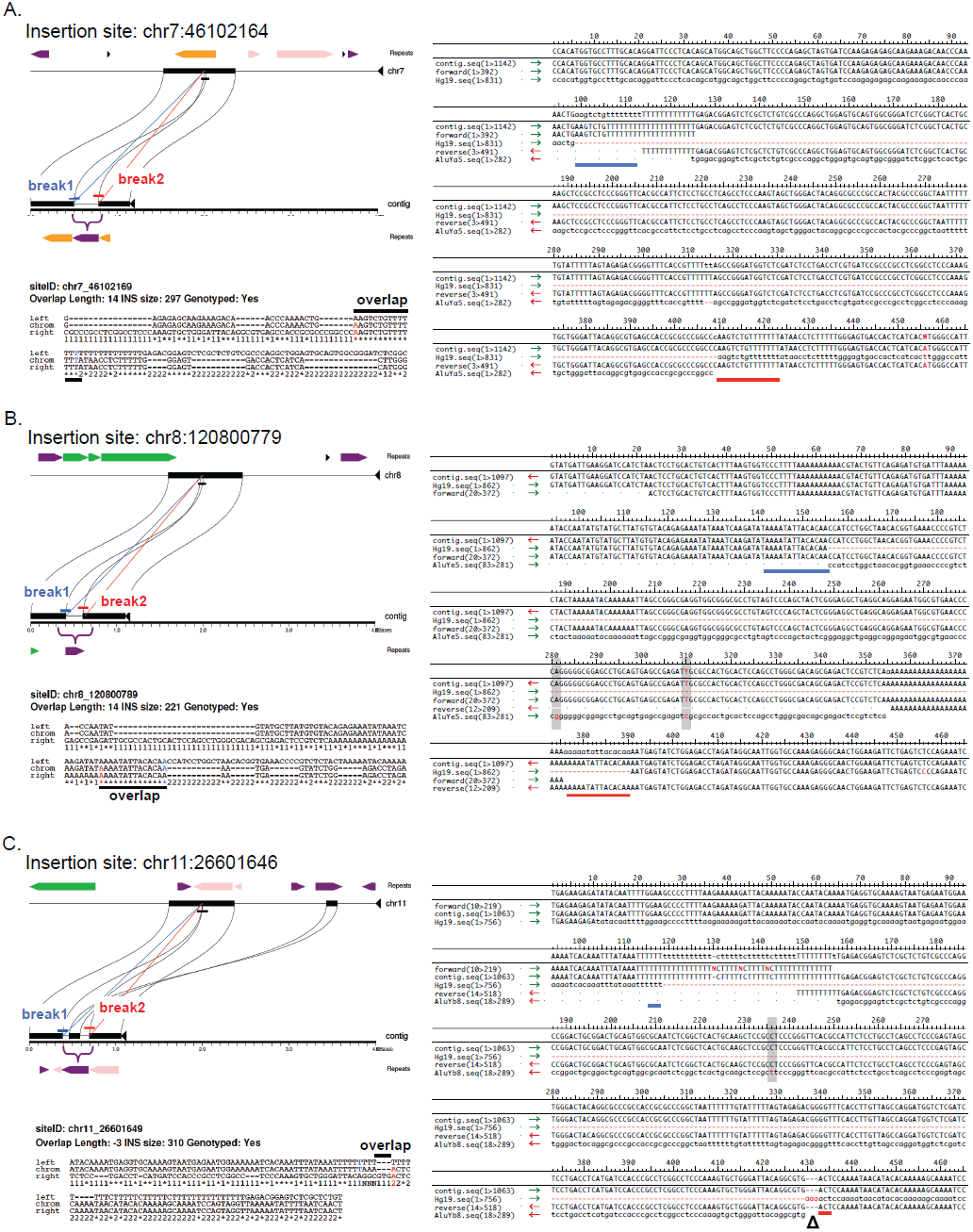
Sequence analysis of assembled non-reference *Alu* insertions. Breakpoints were determined based on alignment the 5’ and 3’ edge of each insertion sequence with the corresponding sequence from the hg19 reference. *Miropeats* annotation, aligned breakpoints, and Sanger sequences are shown for three representative insertions. A. Insertion at chr7:46102164 of 297 bp *Alu* with breakpoint overlap of 14 bp. *Upper left:* Alignment and breakpoint prediction of the assembled contig to hg19. Aligned breaks are shown in blue or red (leftmost or rightmost aligned nucleotides, respectively); the bracket indicates the *Alu* location in the contig relative to hg19. Repetitive elements in the reference and assembled contig are shaded: LINEs, green; SINEs, purple; LTRs, orange; DNA elements, pink. *Lower left:* 3-way alignment of *Alu*-flanking assembled stretches to hg19. A ‘1’ or ‘2’ indicates nucleotides aligned between the assembled contig and hg19 reference upstream or downstream of the *Alu* junction. A ‘*’ indicates identical positions. Terminal nucleotides of the left and right breaks are shaded as above; the black bar shows contig overlap. *Right:* Alignment of the assembled contig with Sanger sequence data to the hg19 empty allele and subfamily consensus for that insertion. Blue and red bars indicate left and right breaks; shading shows assembled and validated base changes from the subfamily consensus. B. Insertion at chr8:120800779 of 221 bp with a 14 bp TSD. C. Insertion at chr11:26601646 of 310 bp with a target site deletion of 3 bp (‘∆’ in the alignment). All breakpoint and alignment indications are as described in panel A.

The presence of each of the 20 randomly selected sites was validated in at least two individuals predicted to have the insertion. Inspection of individual nucleotide alignments confirmed the sequence accuracy of each assembled element (base changes relative to subfamily consensus sequences are shaded in each alignment in Figure S4); however, there were differences observed in breakpoints and flanking sequences for some sites. The randomly selected set of 20 includes 13 insertions deemed suitable for genotyping. The predicted sequence was confirmed for each of these 13 elements. For one site, located at chr6:131113270, the candidate TSD could be extended to include additional sequence that was originally excluded due to ambiguities in alignment gap placement. Seven of the 20 sites had predicted breakpoints within 100 bp of a gap in the CAP3 assembly. Experimental validation resolved the sequence corresponding to the gaps for 6 of these 7 insertions. For one site, at chr6:156541100, the CAP3 assembly gap was within the element poly-A tail and could not be resolved. The predicted breakpoint for 5 of these 7 insertions was confirmed, including 2 sites with additional predicted sequence embedded within the poly-A tail that could not be confirmed.

We also validated a set of 33 calls biased to include insertions predicted to have unusual breakpoint characteristics (*i.e.,* 0-3 bp TSDs, regions of sequence overlap larger than 20 bp, sites with corresponding target site deletions, and insertions with predicted 5’ truncations). We successfully confirmed the presence and predicted nucleotide sequence of each assembled *Alu* element (nucleotide alignments are in Figure S5). Overall, 24/33 insertions had breakpoints that precisely agreed with the corresponding assembly (for example, see Figure 2A). Of the 26 of these 33 sites utilized for *in silico* genotyping 3 were found to have incorrect breakpoints, in each case due to the absence of target site sequence adjacent to the *Alu* poly-A tail in the CAP3 assembly. The 24 fully validated sequences include a full-length element having a 20 bp candidate TSD within a 46 bp region of overlap which likely represents deletion of a pre-existing insertion (chr18:74638702), 3 assembled insertions that were correctly predicted to have short target site deletions relative to the pre-insertion allele ranging in size from 1 to 6 bp (at positions chr6:164161904, chr11:26601646, and chr12:73056650; see also Figures 2B and S5), and several elements with evidence of 5’ truncation (also detailed further below). We were also able to precisely reconstruct insertions that were within other repetitive sequence classes (for example, see Figure 2C).

Finally, we validated each of the 13 assembled elements belonging to the *Alu*S and *Alu*J lineages (Table S3). We confirmed both the variable presence and derived subfamily of the element for each of these sites (detailed alignments are provided in Figure S6). One of the 13 contained an assembly gap near the predicted breakpoint. As anticipated, several insertions had sequence characteristics indicating variable presence caused by non-retrotransposition mechanisms (*i.e.,* involved in encompassing deletions or other endonuclease-independent processes) that were captured in our assembly and experimentally validated; these particular sites are detailed further below.

### Characteristics of assembled insertions

Given the accuracy of our assemblies, we sought to more comprehensively characterize our set of reconstructed *Alu* insertions. Previous studies of full-length polymorphic elements have been mostly limited to insertions from an assembled reference genome (13,79), examination of trace archive data (12), insertions having been captured in relatively long read data (10), or consensus sequence generation based on re-mapping of reads (14). By making use of WGS data in *de novo* assembly, the insertion sequence itself is accurately reconstructed for analysis. Thus, utilizing our assembled contigs, we readily extracted the corresponding 1,614 *Alu* nucleotide sequences and characterized each in terms of subfamily distributions and element properties.

Based on sequence divergence from *Alu* subfamily consensus sequences obtained from RepBase (55), we were able to assign 1,465 (90.7%) of our insertions to one of 30 subfamilies (Table 2). We found 149 elements that were equally diverged from more than one subfamily consensus and could not be conclusively classified. Insertions from *Alu*Y subfamilies made up >99% of all assigned calls, with *Alu*Ya5 and Yb8 collectively representing more than half (62.7%) of the set. This observation was anticipated, given that *Alu*Y insertions have contributed to nearly all *Alu* genomic variation in humans, with *Alu*Ya5 and Yb8 being the most active subfamilies (5,9); and is broadly consistent with the subfamily assignments obtained by Mustafa *et al.* (14). Also as expected, assembled elements derived from the older *Alu*S and *Alu*J lineages were a minority, together representing less than 1% of calls that could be assigned to a subfamily and similar to previous analysis of representative intact polymorphic *Alu* in humans (10–13).

**Table 2.** Classification of assembled *Alu* variants.

| Subfamily | Count | % Total | % Assigned |
| --- | --- | --- | --- |
| ***AluY*** |  |  |  |
| *AluY* | 87 | 5.4% | 6.0% |
| *AluYa1* | 1 | 0.1% | 0.1% |
| *AluYa4* | 85 | 5.3% | 5.9% |
| *AluYa5* | 538 | 33.3% | 37.1% |
| *AluYa8* | 6 | 0.4% | 0.4% |
| *AluYb3a1* | 6 | 0.4% | 0.4% |
| *AluYb8* | 377 | 23.4% | 26.0% |
| *AluYb9* | 50 | 3.1% | 3.4% |
| *AluYc1* | 107 | 6.6% | 7.4% |
| *AluYc2* | 12 | 0.7% | 0.8% |
| *AluYd8* | 9 | 0.6% | 0.6% |
| *AluYe5* | 71 | 4.4% | 4.9% |
| *AluYf1* | 7 | 0.4% | 0.5% |
| *AluYg6* | 50 | 3.1% | 3.4% |
| *AluYh3* | 1 | 0.1% | 0.1% |
| *AluYh7* | 5 | 0.3% | 0.3% |
| *AluYi6* | 17 | 1.1% | 1.2% |
| *AluYi6_4d* | 7 | 0.4% | 0.5% |
| *AluYj4* | 2 | 0.1% | 0.1% |
| *AluYk11* | 4 | 0.2% | 0.3% |
| *AluYk12* | 5 | 0.3% | 0.3% |
| *AluYk13* | 5 | 0.3% | 0.3% |
| ***AluS*** |  |  |  |
| *AluSc* | 1 | 0.1% | 0.1% |
| *AluSc8* | 1 | 0.1% | 0.1% |
| *AluSp* | 1 | 0.1% | 0.1% |
| *AluSq* | 2 | 0.1% | 0.1% |
| *AluSq2* | 1 | 0.1% | 0.1% |
| *AluSx1* | 1 | 0.1% | 0.1% |
| *AluSx3* | 3 | 0.2% | 0.2% |
| *AluSz6* | 1 | 0.1% | 0.1% |
| ***AluJ*** |  |  |  |
| *AluJb* | 2 | 0.1% | 0.1% |
| Unclassified | 149 | 9.2% | 10.3% |
| **Total** | **1614** |  |  |
| **Total Classified** | **1452** |  |  |

Since *Alu*S and *Alu*J are considered to be functionally inactive in humans (5,80), we examined each of the 13 validated sites from each of these families to infer possible mechanisms giving rise to these particular variants (Figures S6 and S7). Several of these insertions had sequence characteristics indicative of a deletion, including extended but non-identical sequence flanking the element that exists as a single copy in the reference (for example, refer to the *AluS*q at chr12:26958660 and *Alu*S×3 at chr17:46617220) or the presence of remnant “scar” sequence indicative of a deletion (*Alu*Sq2 at chr10:68049106). Of the remaining elements, one (*Alu*Sc8, chr6:31296783) is found within the HLA region on chromosome 6 and present on alternative HLA haplotypes, suggesting it has been maintained as polymorphic in the human population (81). Surprisingly, four of the elements have nearly identical matches with *Alu*J or *Alu*S sequences located elsewhere in the genome (Figure S7), two of which also include matching non-*Alu* sequence, suggestive of their presence by non-TPRT mechanisms.

We searched for evidence of each of the *Alu*S and *Alu*J insertions in non-human primates, utilizing reference genome sequences and intersection with polymorphic insertions reported in chimpanzees, gorillas, and orangutans by Hormozdiari *et al.* (67) (also see Methods). We found that 7 of the 13 elements were present in the genomes of Great Apes and other primates, with 3 also reported as polymorphic in Hormozdiari *et al.* (also refer to Table S5). The presence of these shared sites again implies that the variable presence of these elements in humans is not caused by new retrotransposition events, but rather is a consequence of other processes.

### Insertions with 5’ truncation

To assess the length distribution of non-reference *Alu* variants from our call set, we focused on insertions assembled from the *Alu*Ya5 and *Alu*Yb8 subfamilies. We reasoned that analysis of these particular subfamilies should provide the most informative resource for comparison given their representation as the majority of identified variants. We further limited analysis to those *Alu* that were suitable for genotyping, as insertions that do not meet our criteria for genotyping may erroneously appear to be truncated due to an incomplete breakpoint assembly. This resulted in an analysis set of 351 *Alu*Ya5 and 215 *Alu*Yb8 insertions. Based on nucleotide alignments of the assembled insertions against their respective consensus, we examined the collective coverage of assembled elements, per subfamily, in comparison to the nucleotide positions relative to their respective consensus (also refer to the plots in Figure 3A and B).

**Figure 3.**
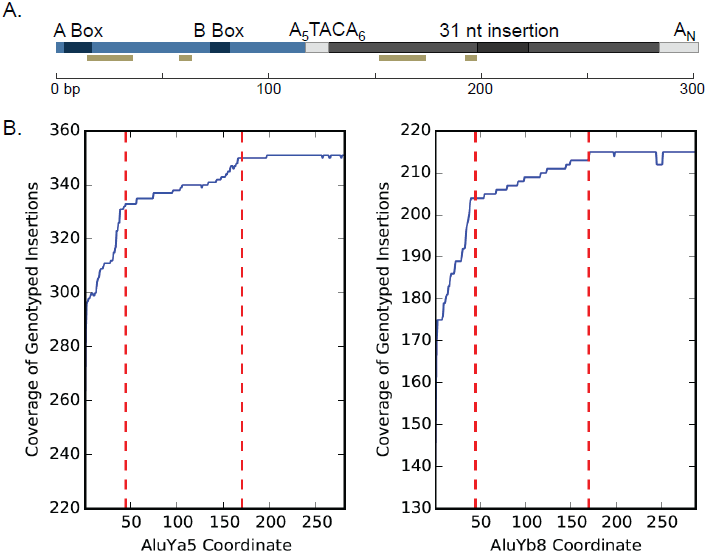
Length distribution of assembled *Alu*Ya5 and *Alu*Yb8 insertions. A. Scaled representation of the *Alu*Ya5 and *Alu*Yb8 consensus and element properties. The *Alu* is comprised of two arms (*left,* blue; *right,* grey) joined by an A-rich region and having a 3’ poly-A tail. The A and B boxes indicate promoter regions. A 31bp insertion distinguishes the arms. *Alu*Yb8 has a 3’ 7 bp insertion relative to Ya5; the sequences are otherwise structurally conserved. The gold bar shows bases within SRP/14 binding sites. B. The size distribution of 351 *Alu*Ya5 and 215 *Alu*Yb8 assembled insertions relative to the respective subfamily consensus. The red dashed lines indicate peaks in truncation near 45 bp and 170 bp. The number of assembled insertions containing an aligned nucleotide is shown against the corresponding position in the consensus.

We observed that 84.9% of *Alu*Ya5 (298/351) and 81.4% *Alu*Yb8 (175/215) variants were full-length, or within at least 5 bp of being full-length, consistent with previous reports of the genome-wide distribution of full-length *Alu* (31,82). Comparing the length distribution of all insertions revealed a detectable minority of 5’ truncations that were present in both subfamilies and exhibited a similar distribution of the apparent truncation point, as is shown in Figure 3B. More specifically, a subset of insertions from either subfamily was truncated ~8–45 bp from the consensus start (9.9% or 35/351 *Alu*Ya5, and 13.4% or 29/215 *Alu*Yb8 insertions), and a second subset was truncated ~55–170 bp from the consensus start (5.1% or 18/351 *Alu*Ya5, and 5% or 11/215 *Alu*Yb8) (Table 3). Besides having apparent 5’ truncation, all but two of these assembled insertions displayed characteristics of standard retrotransposition, including short flanking duplicated sequence and a poly-A tail (insertions at chr13:86166445 and chr11:26601646; also see Table S1). We note the observed distribution is similar to that from the previous analysis of 10,062 reference human *Alu* (as extracted from NCBI build33) (82), and of 1,402 intact polymorphic *Alu* from the then-current dbRIP (31); aspects of both are addressed further in the Discussion.

**Table 3.** Truncation analysis of *Alu* variants.

| Subfamily | Start | Count | % total |
| --- | --- | --- | --- |
| ***AluYa5*** |  |  |  |
|  | 1 to 5 bp | 298 | 84.90% |
|  | 7 to 45 bp | 35 | 9.97% |
|  | 55 to 171 bp | 18 | 5.12% |
| ***AluYb8*** |  |  |  |
|  | 1 to 5 bp | 175 | 81.39% |
|  | 7 to 45 bp | 29 | 13.48% |
|  | 55 to 171 bp | 11 | 5.11% |

L1 and *Alu* insertions that are truncated but otherwise standard (*i.e.,* have poly-A tails and flanking TSDs) are thought to arise from incomplete or premature TPRT (30,31,82,83). One mechanism thought to contribute to 5’ truncations is a microhomology-mediated pairing of nucleotides at the genomic target 5’ end with the nascent *Alu* mRNA, thus promoting the premature completion of TPRT (31), and in turn leaving a detectable signature. We manually examined each three way alignment of the 53 *Alu*Ya5 and 40 *Alu*Yb8 assembled 5’ truncation events for such evidence, specifically searching for nucleotides at the 5’ break that were shared with the respective *Alu* consensus at that position (31). We observed a subset of insertions with detected microhomology, with 40.9% of truncation having 1 bp of matching sequence, and 15.1% of all truncations with ≥ 2 bp shared at the 5’ break (details are summarized in Table S4), though we note limitations of interpreting a single shared nucleotide as a ‘true’ instance of microhomology. Given this observation, the data indicate premature TPRT may account for a proportion of the truncated insertions.

### Insertion breakpoint distribution

We analysed the distribution of assembled insertions based on Gencode v19 annotations (84) relative to simulated placement of insertions at randomly sampled genomic locations, utilizing a position probability matrix (PPM) based on L1 EN preferred nic sites as reported by Gilbert *et al.* 2005 (68) and excluding positions within 500 bp of annotated *Alus* in the reference. Of the 1,614 assembled insertions, 865 (~53.5%) were found within genes, of which 643 (~39.8%) were located within protein coding genes. Although slightly higher than reported in previous analysis (10,11), these values are consistent with our simulated placements (87.5% of simulated insertion sets had ≥ 865 placements within genes, and 82% had ≥ 643 placements within protein coding genes). Just 10 insertions (~0.61% of all calls) were found within exons, all of which were located in untranslated regions and therefore would not be predicted to disrupt coding sequence. This value is lower than that obtained in our simulation (mean of 32 sites with 0 of 200 random simulations having ≤ 10 insertions), indicating potential selection against retrotransposition into exons and other coding sequence, and consistent with previous studies indicating exonic depletion of *Alu* (10,11,34,35).

A total of 708 (~43.8%) of our assembled insertions were located within repetitive sequence. The majority these insertions were found within other retrotransposon-derived elements (459, or ~28.4%, were in LINEs and 124, or ~7.6% in LTRs), and in DNA transposons (69, or ~4.2%); 22 insertions were found in minor or unknown repetitive classes. This distribution is also consistent with that observed in previous survey of non-reference *Alu* insertions (10). Since we excluded any candidate call that was near an annotated *Alu* prior to assembly, no insertion from our callset was recovered within or near any existing *Alu,* though a handful of insertions (33, or ~0.02%) were observed within non-*Alu* SINE classes (*e.g.,* the *Mir,* FLAM, or FRAM groups). These results are broadly consistent with simulated insertions, except that we observe a deficit of insertions within other SINE classes and within satellite sequences (2.5% of simulations resulted in ≤ 33 insertions in SINEs and 0% of simulations resulted in ≤ 2 insertions in satellite sequence). Separate simulations permitting placement near annotated *Alus* in the human reference show that 48% of insertions sampled based on the calculated PPM would be within 500 bp of a reference *Alu* and subsequently excluded from our analysis. However, we note that none of our simulated data sets reached the level of 54% of insertions near other *Alu* elements that is observed in the Chaisson *et al.* data.

### Genotyping

We identified a subset of 1,010 insertions that had both breakpoints at least 100 bp away from an assembly gap that were suitable for *in silico* genotyping using Illumina sequencing reads (Table S6, Figure S8). We compared the inferred genotypes for 11 autosomal sites with PCR-based genotyping across 10 samples, and found a total concordance rate of 99% (109/110) (Figure 4A and B; predicted genotypes are in Table S7 for direct comparison). The only error among the tested calls occurred when the inferred genotype was homozygous for the insertion allele, while PCR genotyping indicates that the site is heterozygous (chr10:19550721; HGDP00476). Finally, we performed a Principle Component Analysis (PCA) of the autosomal genotypes across all 53 samples (73). As expected, individual samples largely cluster together by population with the first PC separating African from non-African samples (Figure 4C). This result further confirms the high accuracy of the inferred genotypes.

**Figure 4.**
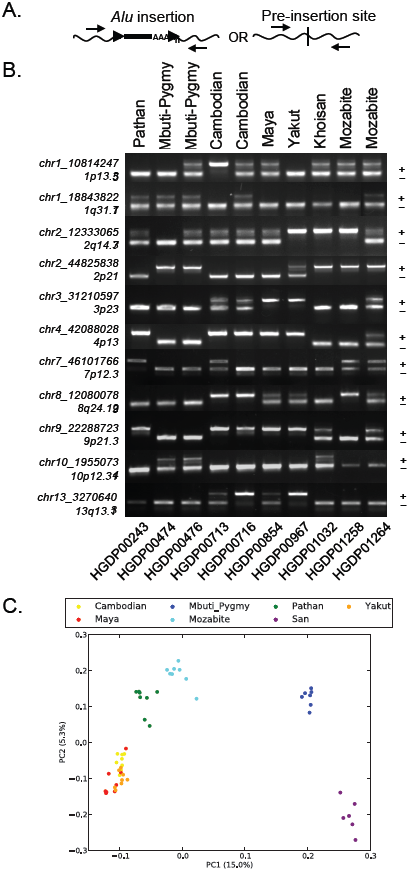
Genotyping of a subset of non-reference *Alu* insertions. Genotype validation was performed for 11 sites across 10 individuals. A. Strategy for primer design and allele detection. A single primer set was used for genotyping each locus, designed to target within 250bp of the assembled insertion coordinates relative to hg19. B. Genotyping from PCR screens and band scoring. Banding patterns supporting the unoccupied or *Alu*-containing allele were assessed following locus-specific PCR; predicted band sizes were estimated by *in silico* PCR analysis and mapped *Alu* coordinates per site. The chromosomal location of each *Alu* is indicated at left. A ‘+’ or ‘-’ shows the relative position of each allele. Sample information is provided for population (above) and for each individual (below). C. Principle Component Analysis PCA was performed on genotype matrix for 1,010 autosomal sites genotypes across 53 populations. A projection of the samples onto the first two Principal Components is shown.

## DISCUSSION

We utilized Illumina WGS paired reads to fully reconstitute a high-specificity set of 1,614 non-reference *Alu* insertions from a subset of 53 genetically diverse individuals in 7 global populations from the HGDP (51,52). Experimental interrogation of a total of 66 sites confirmed the presence and deduced nucleotide sequence of a non-reference *Alu,* at each predicted site. The majority of events appear to result from retrotransposition characterized by apparent TSDs within an expected size range and the presence of poly-A tails, including several insertions with variable 5’ truncations (also see Figures S4 – S6). Mustafa *et al.* also reconstructed polymorphic *Alu* insertions from population samples (14). Our method, which requires assembly across breakpoints of each insertion, recovers a smaller number of variants per sample than Mustafa *et al.,* which created a consensus based on remapping reads for each element. In both studies, *Alu*Ya5, Yb8 and Yb9 account for the majority of calls, but, we recover proportionally more calls classified as Yb8 and fewer Yb9, consistent with the observation that *Alu*Y insertions have contributed to nearly all *Alu* genomic variation in humans, with *Alu*Ya5 and Yb8 being the most active subfamilies (5,9).

We confirmed assembled insertions with aberrant breakpoint characteristics, including breakpoints with deleted sequence relative to the hg19 reference (chr6:164161904, chr11:26601646, and chr12:73056650), as well as insertions for which the TSD was absent (for example at chr17:46617220). For 1,010 insertions that had at least 100 bp of assembled sequence flanking both sides, we obtained a high level of breakpoint accuracy, having perfect agreement at 46 of a total of 51 sites tested (90.2%), including those with predicted aberrant breakpoints and/or 5’ truncation. Analysis of SNPs has demonstrated that improvements in accuracy can be obtained by separating the “discovery” and “genotyping” phases of analysis (85). We therefore performed genotyping by determining genotype-likelihoods based on remapping Illumina read-pairs to the reconstructed reference and alternative haplotypes, achieving an estimated 99% genotype concordance (109 of 110 genotypes analysed).

For each of the 66 validated insertions, comparisons with Sanger sequences of those sites revealed that the correct nucleotide sequence of the *Alu* insertion itself was obtained in assembly. However, a closer comparison of breakpoints at individual sites indicated that elements located near edges of assembled contigs (*e.g.,* excluding the complete predicted TSD length) were more likely to have incompletely assembled breakpoints. Further examination of the individual reads supporting the assembled contig indicated that this was due to aberrant joining or incomplete TSD capture of reads that covered the poly-A tract (for example, refer to trace data from insertions at chr1:102849294, chr12:99227704, chr22:26997608, and chrX:5781742 in the Supplement). An illustrative example of this comes from our assembly of the previously reported Y *Alu* Polymorphic element (YAP) (86) located at chrY:21611993, which contained an incomplete 3’ TSD. Capillary sequencing in sample HGDP00213 revealed the correct 11 bp TSD (5’ AAAGAAATATA), and confirmed the presence of YAP-specific nucleotide markers (at bases 64, 207, 243, and 268 relative to the *Alu*Y8b consensus), as recovered by our CAP3 assembly and consistent with previous reports (Figure S5.33) (86). Even when alleles are fully (and correctly) reconstructed by read assembly, we note that interpretation of the variant may not be clear. The assembled *Alu*S×3 at chr11:35425392 exemplifies this complexity (Figure S6.8). At this site, our breakpoint predictions were inaccurate due to the presence of concomitant variation at this site relative to the hg19 reference, as revealed by sequencing in other individuals without the insertion to better reconstruct the structure of the pre-insertion allele (Figure S9). Given the structure associated with this insertion, we suggest its variable presence is the result of an encompassing deletion. Notably, the CAP3 assembled insertion and additional non-reference proximal genomic sequence was found to be in complete agreement with corresponding Sanger reads, despite the presence of this surrounding structural variation relative to the genome reference.

Our validations of 11 *Alu*S and 2 *Alu*J insertions (Table S3) correctly confirmed their bimorphic presence and high level of divergence from their respective subfamily consensus (Figure S6). Surprisingly, we identified four of these elements (including both assembled *Alu*Jb copies) that had nearly identical matches with *Alu*S or *Alu*J sequences located elsewhere in the genome (Table S3 and Figure S7) implying a common source. As these elements have shared accumulated mismatches from their subfamily consensus in divergences ranging from ~13.2 % to 14.2 %, we suggest these particular elements are likely the consequence of non-TPRT processes. Two of these elements lacked TSDs and/or poly-A tail (*Alu*Jb at chr12:73056650, and *Alu*S×3 at chr17:46617220). The remaining two elements had adjacent non-*Alu* sequence that can be mapped contiguously to other locations within the human reference genome (*Alu*Sp at chr3:110413394 and *Alu*Jb at chr5:172054822). Multiple non-TPRT mechanisms may have resulted in the duplication of these sequences. For example, they may have served as donor sequences utilized in replication template switching (87,88) or in templated double-strand break repair (89) as has been reported for CRISPR/Cas induced lesions (90). Several insertions had sequence characteristics indicative of subsequent deletion in the genome reference sequence, such as the presence of regions of overlap of unusual length (*i.e.,* 21 bp overlap for the *Alu*S×1 located at chr2:161952317, 98 bp for the *Alu*S×3 at chr11:35425492), additional flanking non-*Alu* sequence that does not map to the human reference at that location (refer to alignments for sites chr1:232869263, chr6:5348761, and chr12:26958660 in Figure S6), and/or coincident presence with bimorphic insertions reported in the genomes of Great Apes and other primates (67) (Table S5). In addition to the 7 *AluS* variants classified as likely deletions, we identified 8 sites having stretches of identical candidate TSDs that were at least 15 bp shorter than the corresponding regions of sequence overlap, suggesting that ~1% of the assembled events may actually represent deletions.

Having an assembled, high-specificity call set of non-reference *Alu* variants permitted analysis of element properties. Performing this step was meant to take particular advantage of these data, as existing MEI callers are generally designed to catalogue events detected from read-based signatures within the data. Examination of our assembled insertions suggested the majority of elements exhibited properties consistent with classical retrotransposition, specifically being full length and the presence of a TSD and poly-A tail of variable length. However, our analysis of the length distribution of the reconstructed *Alu*Ya5 and Yb8 insertions also revealed that 93 elements (~16.4%) of this subset had evidence of having been 5’ truncated, despite appearing otherwise standard, indicating insertion by mechanisms involving premature TPRT. We also observed evidence of at least two groups of this subset, respectively truncated <45 bp and <170 bp from the canonical 5’ edge (Table 3 and Figure 3).

These data are consistent with a previous manual curation of 1,402 intact polymorphic *Alu* from dbRIP that characterized full-length elements available at the time (31). In that study the authors identified 115 elements (~8.2%) with apparent 5’ truncations ~8–45 bp from the *Alu* start (~8.2%) and 89 elements had ~55–171 bp truncations (6.3%) (31). The authors proposed a model of microhomology-mediated nucleotide pairing of the 5’ end of the genomic strand with the *Alu* RNA, having observed 41.2% events with nucleotides at the 5’ break shared with the *Alu* consensus at that position. However a single shared base supported the majority of the truncations; considering ≥ 2 bp accounted for 16.7% of their observed events. We searched our own data corresponding to all 5’ truncation events, and observed similar levels of putative microhomology: 15.1% had at least 2 shared bases at the 5’ edge, and 40.9% of insertions shared 1 base; although tentatively considered to represent true cases of microhomology. Another study reported similar instances of *Alu* truncation events (1,005/10,062 or ~10.5%), but found little to no statistical support for base overlap at the 5’ breaks (~29% 1 bp; ~13% ≥ 2 bp) (82). Given that the 5’ *Alu* end is particularly GC rich, this suggests such a ‘mis’-pairing during TPRT would account for a minority of observed truncations. In support, we examined the nick site for truncations with and without putative signatures of microhomology and found no difference in preference, further confirming that both classes contained the canonical L1 ORF2 protein (ORF2p) nick site, 5’ T_4_/A_2_ (the ‘/’ indicating the site of cleavage) (91,92).

We note that secondary structure of the *Alu* RNA itself may drive the non-random distribution of 5’ truncation points. The bases associated with the points of truncation, near ~45 bp and ~170 bp from the *Alu* start are also coincident with the predicted hairpin structure in the folded RNA (9). The *Alu* RNA is reverse transcribed by the L1 encoded ORF2p, which pauses at sites of RNA secondary structure such as poly-purine tracts and stem-loops (93). Additionally, both truncation regions are located directly 3’ to predicted SRP9/14 binding locations (7,94). Although SRP9/14 binding is necessary for efficient retrotransposition, the younger *Alu*S and *Alu*Y subfamilies contain nucleotide substitutions that reduce SRP9/14 binding affinity, suggesting that efficient displacement of bound SRP9/14 is important for the successful propagation of these elements (5,9). This suggests that the characteristic location of 5’ truncations may be a consequence of ORF2p pausing and premature disengaging from the *Alu* RNA during reverse transcription. Regardless, the data indicate premature TPRT may account for a subset of the truncated insertions (21,30,83).

Although our assembled calls are of high quality, our discovery process suffers from the same limitations that are common to other studies utilizing NGS. Because of the variability in coverage across samples, we are likely missing sites present in only one or a small number of the analysed samples therefore biasing our call-set toward common insertions. For example, of the 994 genotyped sites on the autosomes or in the X-par region, we predict no singleton genotypes, 5 sites with an allele count of 2, and 147 sites with an allele count ≤ 10 across all 53 individuals (Figure S8). In addition to discovery sensitivity, the genotypes inferred in samples having lower coverage are expected to be less accurate. By requiring element assembly, we focus on a highly reliable call set that will have reduced sensitivity. To further explore these issues, we compared our approach with *Alu* insertions identified in the CHM1 hydatidiform mole using PacBio reads (53). This analysis highlights trade-offs in sensitivity and specificity that are inherent in any discovery approach and clearly demonstrates the challenges for discovering *Alu* insertions that are coincident with existing *Alu* elements in the reference (Table 1).

The observed distribution of insertion sites distant from reference elements is broadly consistent with expectations based on the known L1 EN nic-site sequence preferences. We note that our simulation approach, based on a 5-bp motif determined from cell-culture insertion assays, assumes statistical independence among the positions at the cleavage site. David *et al.* (37) have also analysed insertion positions of a large set of polymorphic *Alus* with respect to sequence motifs, reaching broadly similar conclusions. Their approach directly considers explicit 6-bp sequences by normalizing the number of discovered insertions with a given nic-site by the frequency of the observed 6-mer in the reference genome. The David *et al.* analysis, which included a method for discovering polymorphic insertions near existing *Alu* elements, did not detect an enrichment of insertions near other *Alus* beyond that predicted by the observed 6-mer sequence preferences. In contrast, the level of coincident insertions observed in the Chaisson *et al.* long read PacBio data exceeds that predicted by our 5-bp model of nic site preference. This may result from preferences that are not modelled by the 5-bp motif as well as difficulty in detecting *Alu* insertions within clusters of reference *Alus* using shorter-read length data (37).

Considering insertions near other *Alu* results in thousands of false calls, necessitating subsequent filtering steps (for example in (36,38,39) but see (37)). When considering only insertions that are distant from existing reference *Alu* insertions, our assembly approach has a moderate sensitivity (77%), and an FDR less than 5%. The true false-call rate of the insertions assembled from Illumina data is likely to be lower. Of the 16 assembled insertions that do not intersect with Chaisson *et al.* calls, three correspond to *Alu* insertions reported by Chaisson *et al.* that are near existing *Alu* elements, but remained in our call set because the initial RetroSeq prediction was at least 500 bp away from a reference element. An additional three calls correspond to more complex variants reported by Chaisson *et al.* involving *Alu* and other repetitive sequence. Counting these six sites reduces the apparent FDR of the assembly-based approach to 3.4%. Since the Chaisson *et al.* call set is itself likely to have missed some calls due to variable coverage and mapping ambiguities, we consider these rates to be merely approximations.

Despite the ability of local-assembly approaches to recover *Alu* insertions with high precision, it is clear that analysis of insertions, and other types of structural variation, within highly repetitive sequence using comparatively short reads remains a major challenge. Analysis of insertion site preferences, population diversity, and insertion rates across individuals and somatic tissues should be cognizant of the severe challenges posed for accurate variant detection in repetitive regions.

## ACKNOWLEDGEMENT

We thank Sarah Emery for technical advice, Ryan Mills for meaningful input and critical reading of the manuscript, John Moran for advice on *Alu* RNA secondary structure insertion characteristics, and Amanda Pendleton for discussion and editorial comments.

## DATA ACCESS

The HGDP *Alu* sequence data from this study is available in the NCBI Database of genomic structural variation (dbVar; http://www.ncbi.nlm.nih.gov/dbvar/) under accession nstd109 and is also available in Supplemental Table S1a. The pipeline for *Alu* assembly and breakpoint analysis is available at https://github.com/KiddLab/insertion-assembly. Scripts for randomly sampling positions based on a PPM are available at https://github.com/KiddLab/random-sample-by-ppm.

## AUTHOR CONTRIBUTION

JHW and JMK designed the study. JHW, AB and NMB performed necessary PCR, sequencing, and sequence-based analysis. JHW and JMK were responsible for all other data analysis. JHW and JMK wrote the paper. All authors have read and approved the final manuscript.

